# Leigh Syndrome-inducing Mutations Affect LRPPRC / SLIRP Complex Formation

**DOI:** 10.1101/2020.04.16.044412

**Authors:** Sandrine Coquille, Stéphane Thore

**Author notes:** **Corresponding Author**, Correspondence should be addressed to Stéphane Thore.

## Abstract

Mitochondria are essential organelles carrying their own genetic information which require specific gene expression processes. The leucine rich pentatricopeptide protein (LRPPRC) and its partner the SRA stem-loop interacting RNA binding protein (SLIRP) form a stable complex implicated in mRNA stability and polyadenylation. LRPPRC/SLIRP complex formation is still poorly characterized. We demonstrate that SLIRP interacts with the N-terminal region of LRPPRC in a RNA independent manner. We further show that the complex is stable in presence of high salt concentration. Point mutation and deletions found in the LRPPRC protein and responsible for the French-Canadian Leigh Syndrome (LSFC) are shown to affect complex formation *in vitro*. Our data are identifying the key region of LRPPRC involved in SLIRP association and showing the direct consequence of various LSFC mutations on the complex formation. Further experiments aiming at deciphering LRPPRC/SLIRP function(s) *in vivo* will benefit from our functional domain characterization.

## INTRODUCTION

LRPPRC is a mammalian RNA-binding protein belonging to the pentatricopeptide repeat (PPR) protein family. This protein family is characterized by a 35 amino acid motif, repeated in tandem, which was predicted to fold into a helix-turn-helix structure (Small and Peeters, 2000). Based on secondary structure and fold prediction algorithm, LRPPRC includes three domains containing canonical PPR motifs interspaced by domains with degenerated helical motifs or disordered regions. LRPPRC is localized in mitochondria but its presence in the nucleus and in the cytoplasm has also been reported (Mili and Piñol-Roma, 2003; Topisirovic *et al*., 2009). LRPPRC is a fairly large protein with multiple functions and whose mechanism of action is yet poorly characterized. Up to now, several studies have demonstrated that LRPPRC can form different protein complexes with functions linked to posttranscriptional gene regulation in mitochondria but also cytoskeleton organization, vesicular trafficking or apoptosis (Liu *et al*., 2002; Gohil *et al*., 2010; Michaud *et al*., 2011).

LRPPRC forms a complex with the protein SLIRP (SRA stem-loop interacting RNA binding protein) in mitochondria, which binds all mRNA species in coding regions without apparent sequence specificity (Sasarman et al., 2010). The LRPPRC/SLIRP complex is necessary to maintain and stabilize a pool of non-translated mRNA (Ruzzenente et al., 2012). It prevents mRNA degradation mediated by PNPase/SUV3 helicase and promotes mRNA polyadenylation by the mitochondrial poly(A) polymerase (MTPAP; Chujo et al., 2012). Although PPR proteins have often pre-determined RNA binding specificity (Barkan et al., 2012; Takenaka *et al*., 2013; Yagi et al., 2013; Yin et al., 2013), the regions of LRPPRC predicted to fold into canonical PPR motifs do not seem to share this property.

A single missense mutation in the LRPPRC gene leading to the substitution of Alanine 354 by a Valine is responsible for the French Canadian type of Leigh Syndrome (LSFC; Mootha et al., 2003). This neurodegenerative disease is characterized by a tissue-specific deficiency in cytochrome C oxidase (COX) leading to a decrease of ATP production by oxidative phosphorylation in mitochondria (Merante et al., 1993). It was shown that the mutated LRPPRC protein presents a reduced steady-state level resulting in mitochondrial mRNA destabilization, especially the COX mRNA (Xu et al., 2004). Recent studies performed in a more diverse population have also shown that others mutations (corresponding notably to the deletions of Valine 866, Lysine 909 and the region from Arginine 1276 to Lysine 1300) can occur within the LRPPRC gene with similar functional consequences (Olahova et al., 2015).

The present study is focused on the *in vitro* biochemical characterization of the LRPPRC/SLIRP complex. We produced the LRPPRC/SLIRP complex in quantity sufficient for biochemical studies. Based on the LRPPRC structure prediction, we initiated a series of LRPPRC truncations to determine that the N-terminal region of LRPPRC is responsible for the binding to SLIRP. We further demonstrated that LSFC mutations found in LRPPRC apparently reduce its capacity to form a stable complex with the protein SLIRP. These studies will help the biochemical studies aiming at understanding the precise mechanism(s) allowing the LRPPRC/SLIRP complex to regulate mitochondrial mRNA half-life, polyadenylation and translation.

## MATERIALS AND METHODS

### Plasmid constructions and cloning

The human cDNA clones of LRPPRC and SLIRP were provided by OriGene (product numbers: SC120536 and SC108657). For each construct described in this study, the region of interest was amplified by PCR using the primers listed in Table 1. The resulting amplicons and expression vectors (derived from pET42) were digested using the BamHI and NcoI restriction sites, ligated and transformed into *E. coli* strain XL1 competent cells. The pET42 derived vectors were designed to express the LRPPRC constructs fused to N-terminal His9 tag and the SLIRP construct fused to N-terminal GST/His6 tag. All constructs were confirmed by DNA sequencing.

**Table 1.**
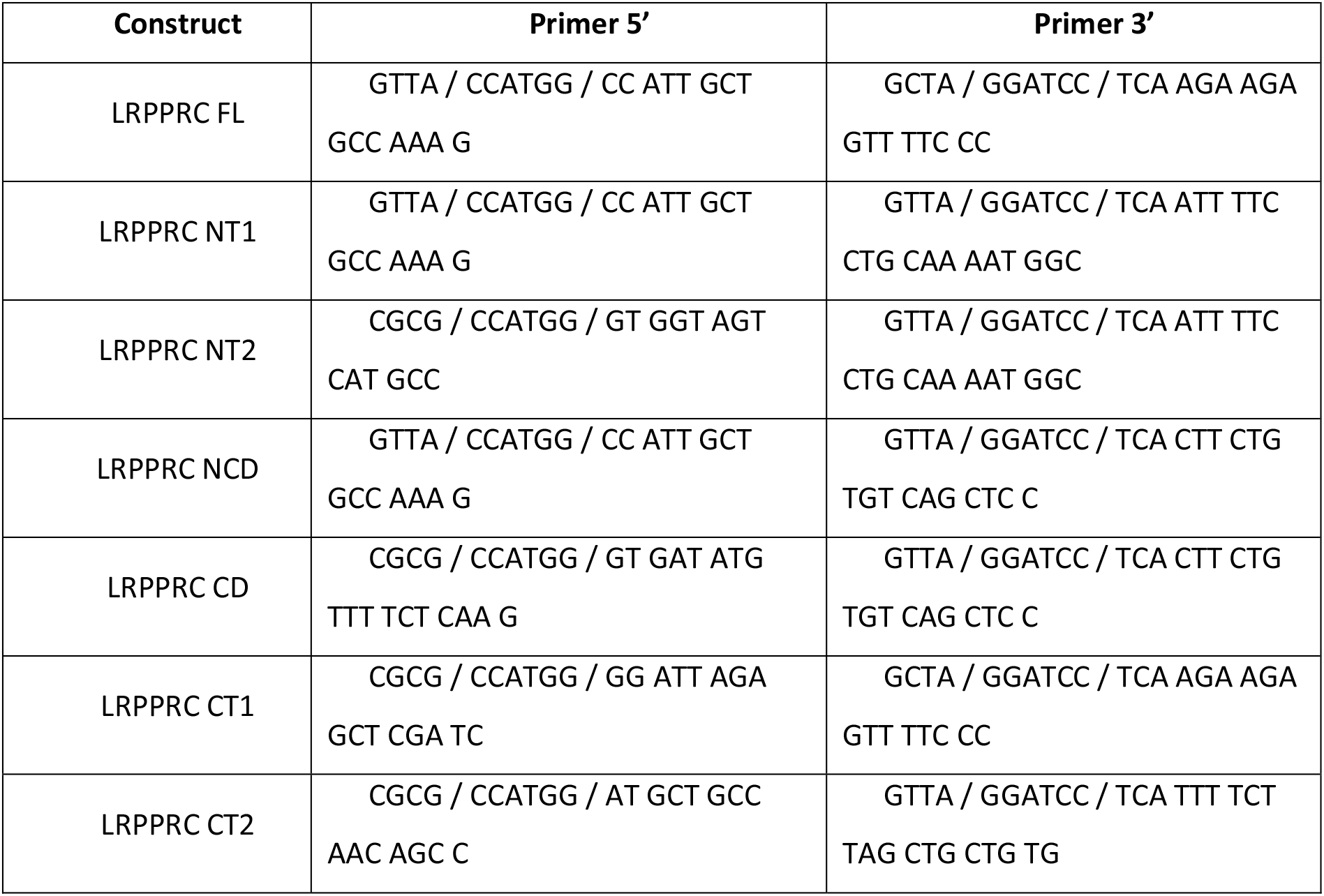
Primers used for LRPPRC cloning.

### Mutagenesis

The LRPPRC A354V, delV866, delK909 and delR1276_K1300 mutations were introduced in the corresponding template vector by site-directed mutagenesis, using the Quick-Change mutagenesis kit (Stratagene) according to manufacturer’s instructions. The primers used for the mutations are shown in Table 2. The constructs were verified by DNA sequencing.

**Table 2.**
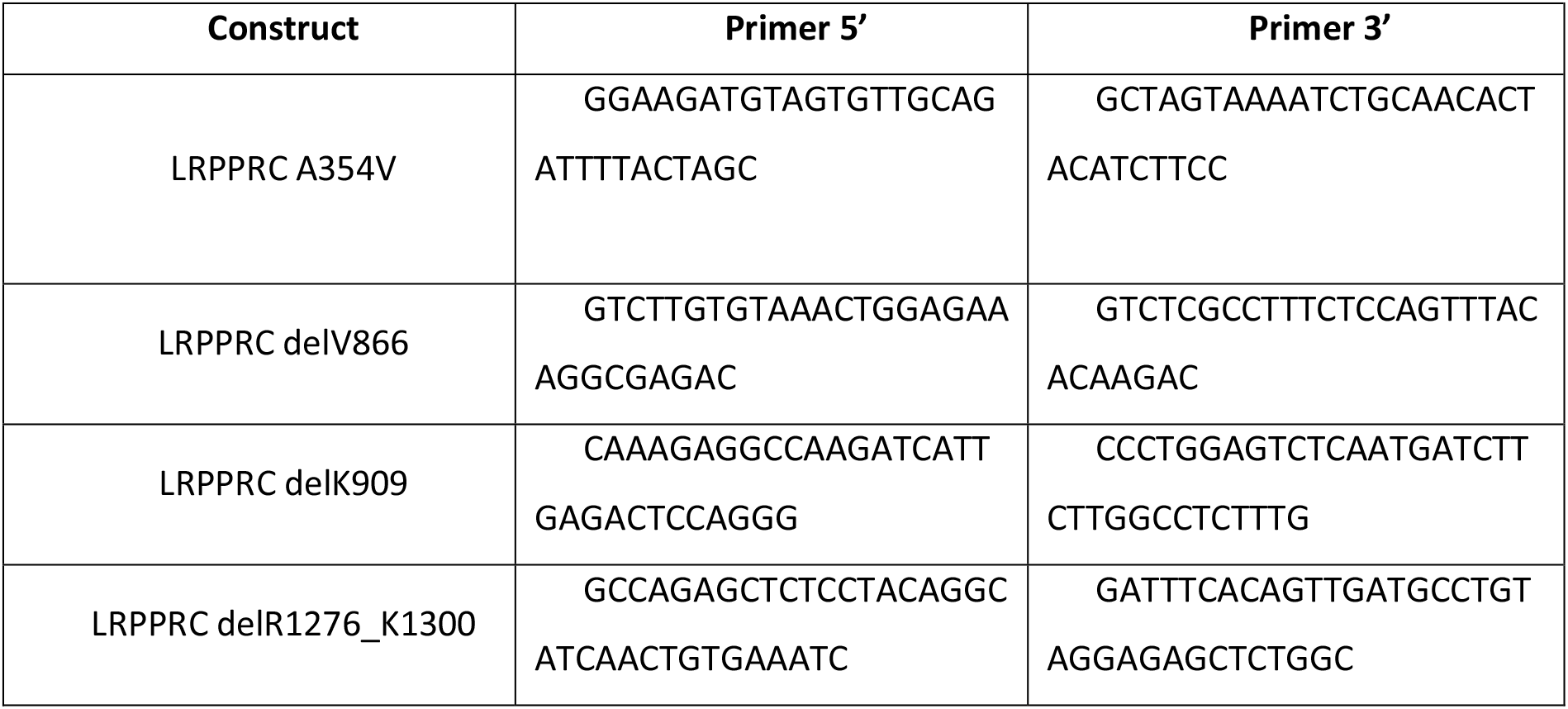
Primers used for LRPPRC site-directed mutagenesis.

### Expression and pull-down experiments

LRPPRC and SLIRP constructs were transformed simultaneously into E. coli strain BL21(DE3) competent cells for protein co-expression. Transformed cells were grown in LB media at 18°C overnight following induction with 0.25 mM isopropyl-β-D-thiogalactopyranoside (IPTG). After harvesting by centrifugation at 5000 rpm, the cell pellets were resuspended in lysis buffer (20 mM Tris-HCl pH 7.5, 150 mM to 2 M NaCl according to the different experiments, 5 mM β-mercaptoethanol) supplemented with DNase 1 μg/ml, Lysozyme 1 μg/ml and protease inhibitors (PhenylMethylSulfonyl Fluoride (PMSF) 1 mM, leupeptin 1 μg/ml, and pepstatin 2 μg/ml).

Cells were lysed using an emulsiflex system (AVESTIN) and cleared by centrifugation at 15000 rpm for 30 minutes at 4°C. The soluble fraction was then loaded on a Protino^®^ GST/4B column 1 ml (Macherey-Nagel) for pull-down experiments. For each experiment, the column was washed with 10 column volumes of lysis buffer before the elution step using the following buffer: 20 mM Tris-HCl pH 7.5, 150 mM (up to 2 M NaCl according to the different experiments), 20 mM reduced Glutathione, 5 mM β-mercaptoethanol. Samples of the different fractions (total, soluble, flow through, washes and elution) were collected and analyzed on 12% SDS-PAGE gel.

### LRPPRC secondary structure prediction

The program Phyre (Kelley et al. 2015) was used to predict the overall 3D structure of the various fragments of LRPPRC using the intensive modelling mode. Figure panels of LRPPRC structure prediction were prepared with the software PyMOL (The PyMOL Molecular Graphics System, Version 1.7.4 Schrödinger, LLC.).

## RESULTS

### LRPPRC/SLIRP interaction domains

LRPPRC and SLIRP proteins appear generally to be co-regulated *in vivo* (Baughman et al., 2009; Chujo et al., 2012; Sasarman et al., 2010; Sasarman et al., 2015; Ruzzenente et al., 2012). Indeed, if the level of one protein decreases, the other one will also be reduced. This clearly indicates that both proteins stabilize each other. However, PPR and RRM domains are mostly known for mediating protein/RNA interaction rather than protein/protein interaction with a few exceptions particularly for the RRM domains (Fribourg et al., 2003).

To determine which region(s) of LRPPRC was(were) responsible for mediating the interaction with SLIRP, we first generated a series of LRPPRC truncations (Fig. 1A). To define the protein borders, we used secondary structure prediction server to delimit the folded domains of the LRPPRC protein. This analysis revealed that LRPPRC could be subdivided into a N-terminal mitochondrial targeting sequence followed by 3 larger regions containing PPR motifs separated by respectively 59, 142 and 33 residues. These inter-domains linkers are predicted to contain helical repeats but apparently not of the PPR type (Cheng et al., 2016). We then co-express these various LRPPRC truncations with a GST-tag version of the SLIRP protein and performed pull-down experiments using the GST domain (Fig. 1B).

**Figure 1.**
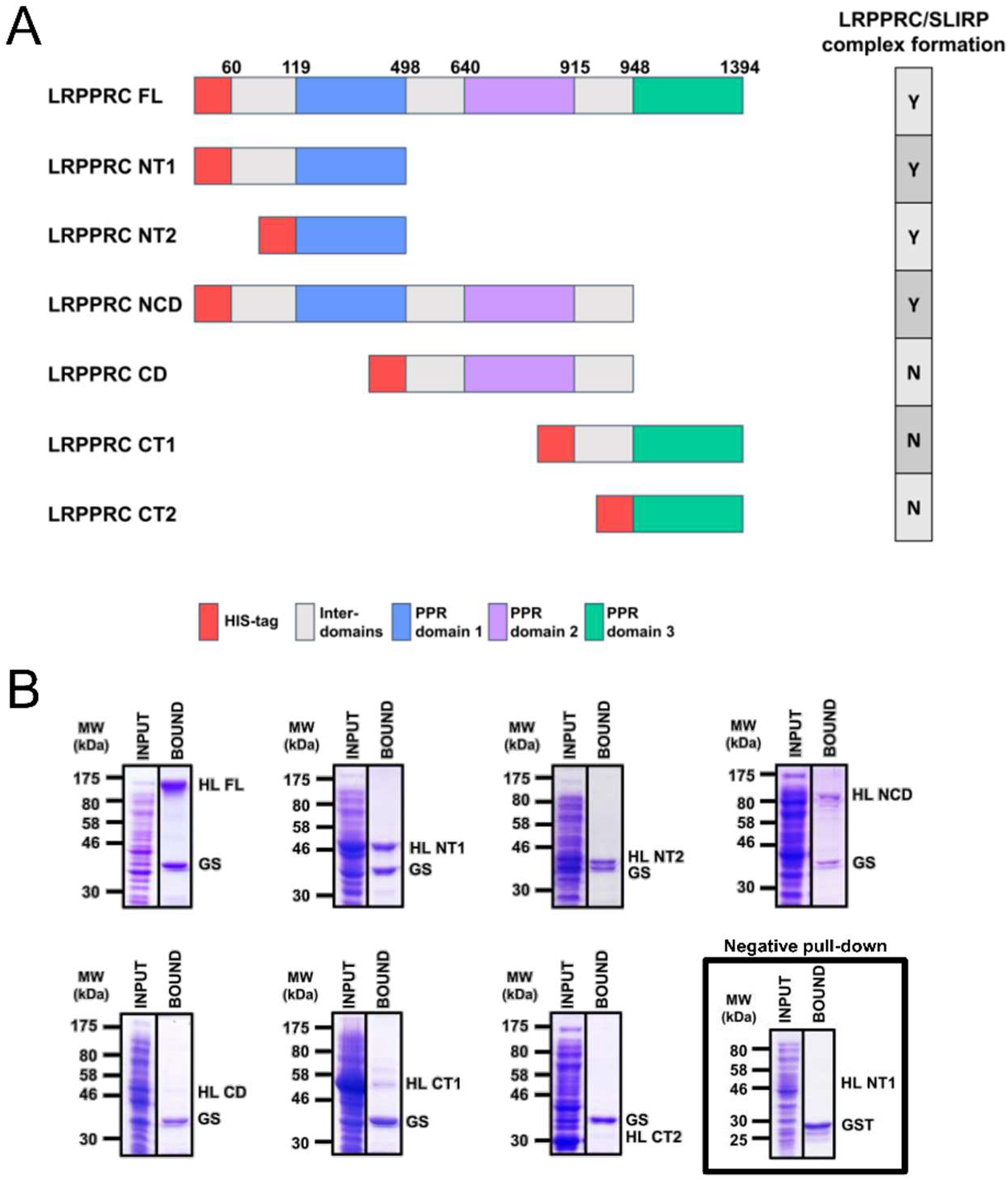
Identification of the minimal region of LRPPRC required to form a stable interaction with SLIRP. A) Schematic view of the LRPPRC proteins with domain annotations. Using secondary structure prediction server, LRPPRC was divided in several fragments. Three domains are formed of PPR motifs while three inter-domains present helical repeats of a different type. Each domain is colored differently. We generated a series of constructs encompassing one or several of these annotated regions. On the right of the diagrams, a table summarizes the complex formation observed between each LRPPRC construct and the GST-tagged SLIRP protein (Y for yes and N for no). B) Affinity purification of the LRPPRC constructs which were co-expressed with a GST-tag version of its partner SLIRP. GST pull-down were performed as described in the material and method section and demonstrated that the N-terminal fragment NT2 is sufficient to observe a stable complex. Input and bound fractions are shown. HL FL stands for His-tag LRPPRC full length; HL NT1 for His-tag N-terminal fragment 1; HL NT2: His-tag N-terminal fragment 2; HL NCD: His-tag N-terminal and central domain; HL CD: His-tag central domain; HL CT1: His-tag C-terminal fragment CT1; HL CT2: His-tag C-terminal fragment 2 and GS: GST-tag SLIRP.

Based on these experiments, we could identify the fragment of LRPPRC containing the SLIRP binding capacity. To exclude that some of the truncations which did not bind SLIRP in our pull-down experiments were in fact unstable or insoluble, we performed affinity purification using Histidine chromatography on these truncations (all the C-terminal fragments of LRPPRC) in order to verify that they were soluble. All these truncations could be purified to homogeneity (data not shown). Thus, the fact that complexes are not formed between SLIRP and the C-terminal parts of LRPPRC is not due to the insolubility of these fragments but seems due to their incapacity to bind SLIRP. We also controlled that the GST tag did not interfere with the complex formation between LRPPRC and SLIRP, i.e. that the different LRPPRC constructs bind SLIRP directly and not through the GST tag attached to SLIRP. For that, we co-expressed the GST tag alone with the His-LRPPRC NT1 construct. By pull-down experiment, we observed no binding between these two constructs confirming the direct association between LRPPRC and SLIRP (Fig. 1B).

*In fine*, LRPPRC sequence could be reduced to the fragment encompassing residues 119 to 498, which correspond to the first annotated PPR region. The identified domain of 379 residues is predicted to fold into 11 consecutive PPR motifs without any obvious domains capable of binding the RRM protein SLIRP. It is therefore likely that structural data will be appreciable to fully understand how these two proteins interact with each other. In all these pull-down experiments, the intensities of the Coomassie-stained bands corresponding to our proteins of interest are quite similar suggesting that the purified complex is likely stoichiometric (Fig. 1B).

Next, we decided to test the strength of the LRPPRC/SLIRP interaction to evaluate whether the absence of the native RNA targets or a high concentration of salt may affect the stability of the complex. As the bound complex was apparently stoichiometric, we increased and decreased salt concentration present during the affinity purification to directly assess whether the interaction surface was likely limited to electrostatic contacts or if it could also contain hydrophobic pockets, potentially less sensitive to the presence of high salt concentration. We thus tested the effect of sodium chloride salt concentration from 150mM up to 2M on the stability of the complex formed between our GST-SLIRP construct and the best LRPPRC binder, i.e. the HL NT1. It is reasonable to assume that 2M salt is the highest concentration of salt to test as protein will start to unfold at higher concentration, unless the protein belongs to a halophilic organism. In our experiment, we can observe that the band corresponding to the HL NT1 construct slightly increases when the salt concentration increases from 150 mM to 500 mM (Fig. 2, lanes 1 and 6). Then, the LRPPRC NT1 band was unchanged between 500 mM and 1 M (Fig. 2, lanes 5 to 8). Finally, the association is lost at a salt concentration of 2 M, likely because proteins start to unfold as mentioned above (Fig. 2, lanes 9-10). Such a stable interaction can be explained if the interaction surface between these two proteins is either very large or is formed through hydrophobic contact enhancing the weak salting-out effect due to the increased in salt concentration.

**Figure 2.**
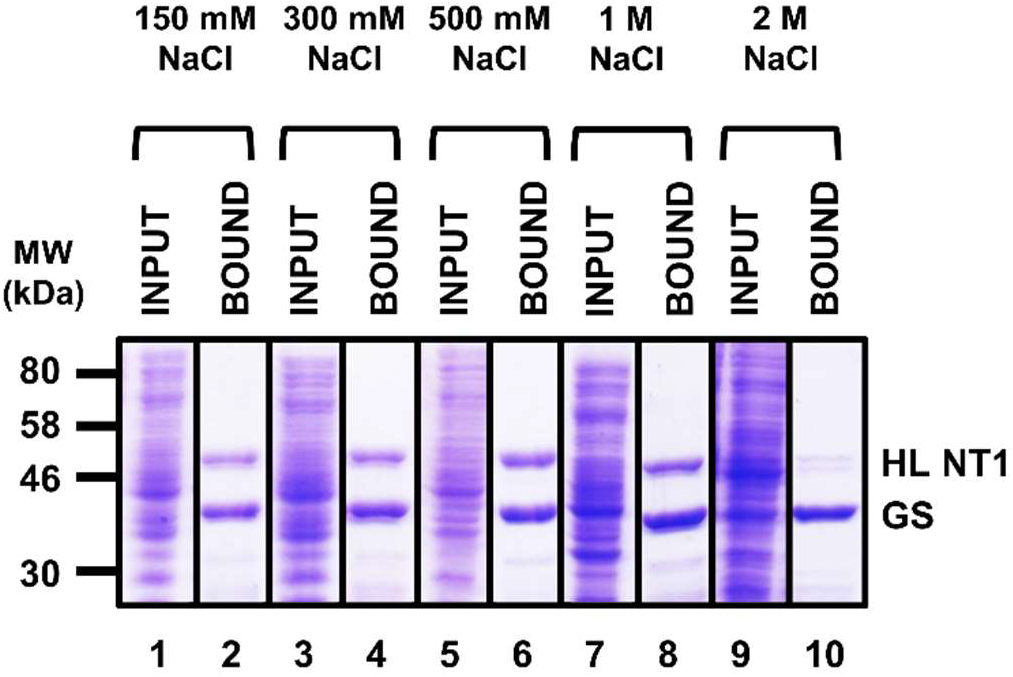
LRPPRC/SLIRP complex stability is almost independent of salt concentration. Cells were lysed in identical buffer with the exception to their respective salt concentrations. Washes were also performed with the same salt concentrations. The bound proteins were finally eluted with 20 mM of reduced glutathione and loaded onto a 12% SDS-PAGE. HL NT1 and GS are as in Figure 1.

The LRPPRC protein is known to be the key factor which the deficiency of leads to the Leigh Syndrome, i.e. a mitochondrial disease (Leigh 1951; Willems *et al*., 1977). A major mutation observed in patient cells presenting this disease maps to the chromosome 2p16-21 in the gene encoding the LRPPRC protein (Mootha *et al*., 2003). This missense mutation, A354V, is located in the N-terminal part of the LRPPRC protein, which we have identified as being essential for the binding to SLIRP in the present study. Furthermore, this mutation is known to affect protein stability since patient with this disease have low level of LRPPRC *in* vivo. (Xu *et al*., 2004; Sasarman *et al*., 2010). Interestingly, SLIRP level is also low in the mitochondria of these patient and LRPPRC is known to affect SLIRP stability, possibly linking default of association with presence of the mutation. We thus attempted to pull-down the NT1 truncation containing the A354V mutation using the GST-SLIRP protein as bait (compare Fig. 1B and Fig. 3A, lanes 3-4). No complex could be observed even when we varied the salt concentration (Fig. 3A, lanes 1-2 and 5-6). These data strongly suggest that the A354V mutation acts negatively on the stability of the LRPPRC/SLIRP complex, thus leading to the degradation of both LRPPRC and SLIRP proteins *in vivo*.

**Figure 3.**
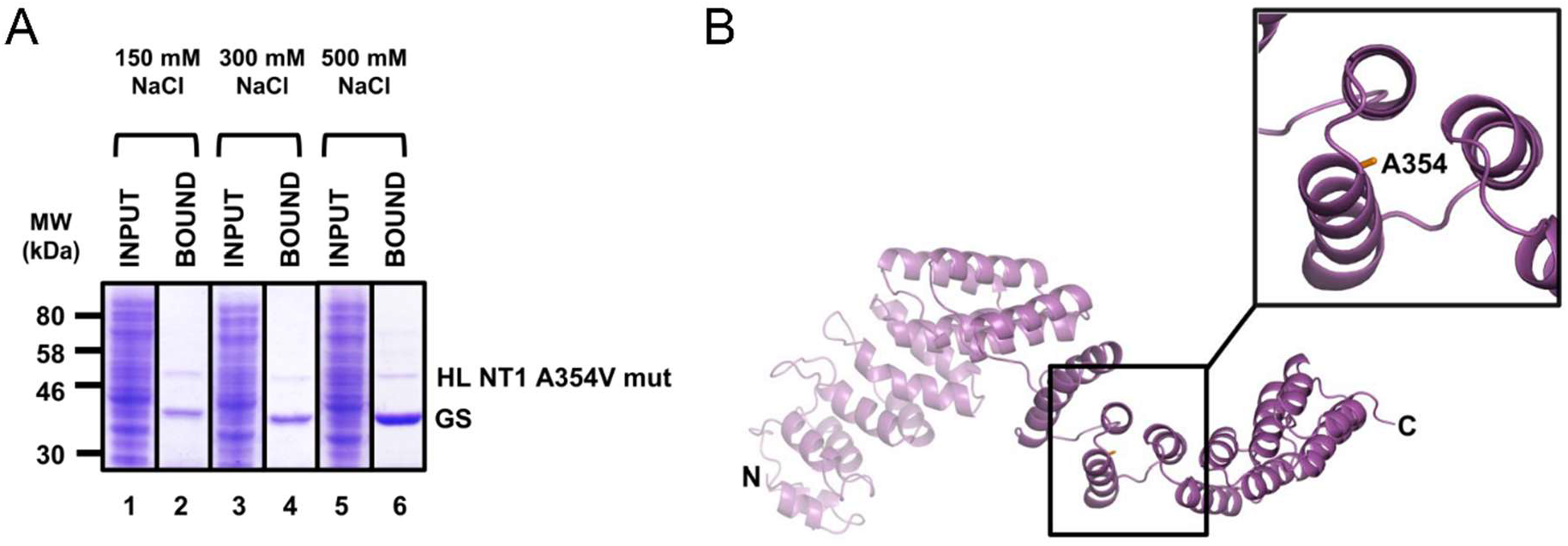
Effect of the LSFC-inducing A354V mutation on LRPPRC/SLIRP complex formation. A) The NT1 construct containing the A354V mutation was co-expressed with the GST-SLIRP protein and pull-down performed. While GST-SLIRP was binding the GST column, the upper band corresponding to the NT1 A354V construct was hardly retained, independently of the salt concentration, indicating that mutated LRPPRC protein does not bind stably to SLIRP anymore. HL NT1 A354V mut stands for His-tag N-terminal fragment 1 carrying the A354V substitution and GS: GST-tag SLIRP protein. B) LRPPRC NT1 structure prediction. The NT1 fragment sequence (residues 60-498) was modeled by the webserver Phyre. The best structural prediction is shown here and was built onto the atomic structure of the PPR10 protein from maize (PDB ID 4M57). The helices surrounding the A354V mutation are shown in the inset. The Alanine residue 354 is shown as stick and colored in orange.

Recently, another group of mutations mapping in the LRPPRC gene have been identified in patients originating outside of the French-Canadian population, i.e. outside of the population in which the founder variant of Leigh Syndrome was first discovered (Oláhová et al., 2015). Two of these mutations are in frame deletions of a triplet of bases in LRPPRC gene, resulting in the deletions of Valine 866 and Lysine 909 in LRPPRC protein respectively (Oláhová et al., 2015). These mutations are located in the central PPR domain of LRPPRC protein. The third mutation is an in frame deletion of 75 bases, leading to the deletion of 25 residues in the translated LRPPRC protein (from Arginine 1276 to Lysine 1300; Oláhová et al., 2015). This third genetic alteration is located in the third PPR domain of LRPPRC. By site-directed mutagenesis, we introduced these mutations in the full-length construct of LRPPRC and we performed pull-down experiments using the GST-tag SLIRP protein as a bait.

We observed that the LRPPRC/SLIRP complex formation is maintained in the cases of LRPPRC single deletions delV866 and delK909, although slightly less when compared with the WT protein (compare Fig. 4A, lanes 2, 4 and 6). Based on the structural model prediction obtained for these LRPPRC domains, we can see that the Valine 866 is located in the hydrophobic core of a PPR repeat and the Lysine 909 is positioned in the second helix of a PPR motif but directed toward the surface (Fig. 4B). Single deletion of these amino acids might thus affect the local conformation of the protein, which could lead to a partial destabilization of the protein compared to the WT one and explain the slight decrease observed in the complex stoichiometry. However, it is unlikely to abolish the complex formation because of the long distance between the presently identified SLIRP binding site and the mutation sites on LRPPRC. Strikingly, the interaction between SLIRP and the LRPPRC carrying the R1276-K1300 deletion seems much weaker or very noisy (Fig. 4A, lane 8). In this case, we rather observe a large number of bands in the pull-down experiments, which are indicating that GST-SLIRP binds to a protein of a variable length. Differently, the deletion from the Arginine 1276 to the Lysine 1300 affects three PPR motifs in LRPPRC C-terminal region according to the structural prediction (Fig. 4C). Indeed, one PPR motif is amputated of the second helix and the two surrounding ones will not be able to interact properly. We hypothesize that the destabilization of the LRPPRC protein C-terminal region is sufficient to partially unfold the protein, thus rendering it sensitive to protein degradation during its production. Instead of observing a single protein band for the partner of SLIRP, we now observe multiple forms of LRPPRC forming a kind of “ladder” in the SDS-PAGE gel (Fig. 4A, lane 8).

**Figure 4.**
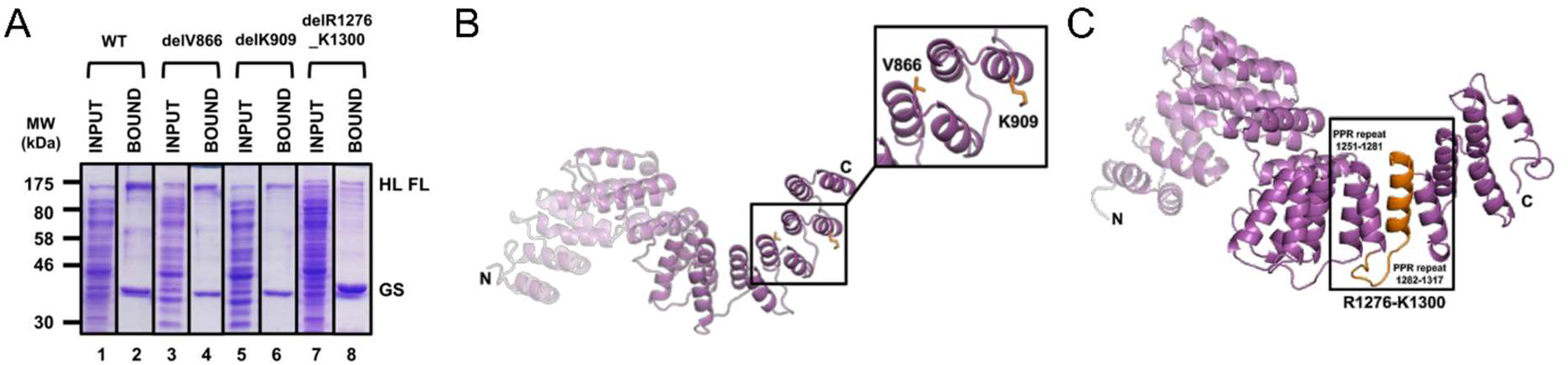
Effect of minor Leigh Syndrome mutations on LRPPRC/SLIRP complex formation and protein stability. A) Pull-down experiments between the full-length LRPPRC constructs containing the different deletions and the GST-SLIRP protein. The constructs with delV866 and delK909 do not impair the LRPPRC/SLIRP complex formation while the construct with delR1276_K1300 appears degraded although it is still pull-downed by GST-SLIRP. WT, delV866, delK909 and delR1276_K1300 stand for wild type LRPPRC, LRPPRC deleted of residue V866, of residue K909 and of the fragment between residues R1276 and K1300 respectively. GS is as figure 1A. B) and C) Structure prediction of second and third PPR fragments of LRPPRC, residues 504 to 948 and 915 to 1394 respectively. The CD and the CT domain sequences were modeled as in figure 3. The mutated amino acids V866 and K909 or the deletion from R1276 to K1300 are highlighted in orange.

Also in these deletion cases, the absence of binding to SLIRP or the partial destabilization of LRPPRC would have for result some defects in mitochondrial mRNA processing possibly leading to Leigh Syndrome.

## DISCUSSION

LRPPRC was originally identified in a genome-wide association screen to map LFSC mutations. In fact, a single missense mutation (corresponding to the A354V substitution in the LRPPRC protein) was shown to be present in various patients DNA (Mootha *et al*., 2003). *In vivo* data have demonstrated that LRPPRC protein level is reduced in mitochondria of patient suffering from the LSFC disease (Xu *et al*., 2004). More recently, the protein SLIRP was shown to be the natural partner of LRPPRC and that protein levels were influenced by each other presence. Although direct interaction between LRPPRC and SLIRP was proposed, it was never formerly shown *in vitro* in the absence of their natural RNA targets. We have filled this gap as we have been able to reconstitute LRPPRC/SLIRP complex *in vitro* in absence of over-expressed RNA molecules and in the absence of RNase inhibitors. Moreover, we also show that SLIRP interacts with the N-terminal region of LRPPRC corresponding to the first block of identified PPR motifs likely through a hydrophobic interface as the complex stability is not affected by increasing salt concentrations.

Our data showed that the C-terminal region of LRPPRC which was previously shown to bind RNA, is unlikely impaired in its function by the presence of SLIRP bound to the N-terminal region of LRPPRC (Mili and Piñol-Roma, 2003). This means that the LRPPRC/SLIRP complex may have multiple RNA binding surfaces formed on one hand by the C-terminal PPR domains of LRPPRC, and, on the other hand, by the RNA binding area present in the RRM of SLIRP. Obviously, this remark will only be valid if the RNA binding area present in SLIRP is not mediating protein/protein interaction with LRPPRC.

We also demonstrate that the missense mutation at the 354 position is abolishing *in vitro* the interaction between the identified N-terminal fragment corresponding to residues 119 to 498 and the protein SLIRP.

The region required for the interaction contains multiple repetitions of PPR motifs. Such domains are known to fold as pair of helices packing again each other and forming long solenoid with positive supertwist (Small and Peters 2000; Aubourg *et al*., 2000; Ringel *et al*., 2011; Howard *et al*., 2012). Moreover, recent structural and biochemical investigation using engineered PPR proteins demonstrated that the relative orientation between each pair of helices was governed by a hydrophobic cage buried between those helices (Coquille et al., 2014, Gully et al., 2015). Interestingly, the alanine to valine substitution appears to map into this hydrophobic region based on the structural model prediction (Fig. 3B). One possible hypothesis to explain the default of interaction could be that the A354V mutation induces a change in the concave or the convex surface of LRPPRC which, in turns, prevents the stable association with the protein SLIRP. Absence of binding would lead to LRPPRC degradation and subsequent defect in mRNA processing *in vivo*. Further experiments are required to determine the influence of this mutation on the capacity of individual LRPPRC or SLIRP proteins to associate specifically with mitochondrial RNAs and subsequently to test whether the RNA half-lives are also affected.

Last, we expressed and tested the association between SLIRP and three recently discovered short deletions in LRPPRC proteins which have been found in DNA from patients affected by LS. At least, two of these deletions do not affect significantly the interaction with SLIRP, which was expected since they are located in the second fragment of PPR motifs present in LRPPRC. The last construct that we tested contains a large deletion possibly affecting three PPR motifs from the C-terminal region of LRPPRC. In this case, it is difficult to fully exclude that LRPPRC/SLIRP association is unaffected as the produced LRPPRC protein is apparently highly unstable and tends to degrade during the pull-down experiments. In conclusion, our data define the interacting region between the major mitochondrial RNA processing factor LRPPRC and its natural partner the SLIRP protein. It will be exciting to see if their association relies on the RNA binding surface of the RRM or whether an alternative area of SLIRP is used to form such a salt-stable complex. We also demonstrate that the LSFC mutation found in LRPPRC prevents a stable complex from being formed *in vitro*, and so likely *in vivo*. Finally, we also show that newly identified mutations of the LRPPRC protein found to cause LS are not affecting the association with SLIRP and so are likely to modify the capacity of LRPPRC/SLIRP to interact with other partners or RNAs. Further experiments will be required to fully apprehend the precise defect(s) occurring in these cases.

## AUTHOR INFORMATION

### Author Contributions

The manuscript was written through contributions of all authors. All authors have given approval to the final version of the manuscript.

### Funding Sources

This work was financially supported by grants from the “Ligue Genevoise contre le Cancer” (n°1011) and from the Worldwide Cancer Research (n°14-0346).

### Notes

The authors declare no competing financial interest.

## ACKNOWLEDGMENT

We wish to thank the past members of the laboratory. We also would like to acknowledge the Molecular Biology department of the University of Geneva to facilitate the use of laboratory space and equipment. We are grateful to the Ligue Genevoise contre le Cancer (n°1011) for the salary support and to the Worldwide Cancer Research for facilitating the use of this financial support. The laboratory was further supported by grants from the Swiss National Science foundation (n°31003A_140924 and 31003A_124909).

## Notes

### Competing Interest Statement

The authors have declared no competing interest.

